# CRISPR-CISH: An *in situ* chromogenic DNA repeat detection system for research and life science education

**DOI:** 10.1101/2025.02.20.639243

**Authors:** Bhanu Prakash Potlapalli, Fabian Dassau, Jörg Fuchs, Deboprio Roy Sushmoy, Andreas Houben

**Author notes:** Corresponding authors: (AH) (BPP).

## Abstract

*In situ* hybridization is a technique to visualize specific DNA sequences within nuclei and chromosomes. Various DNA *in situ* fluorescent labeling methods have been developed, which typically involve global DNA denaturation prior to the probe hybridization and often require fluorescence microscopes for visualization. Here, we report the development of a CRISPR/dCas9-mediated chromogenic *in situ* DNA detection (CRISPR-CISH) method that combines chromogenic signal detection with CRISPR imaging. This non-fluorescent approach uses 3’ biotin-labeled tracrRNA and target-specific crRNA to form mature gRNA, which activates dCas9 to bind to target sequences. The subsequent application of streptavidin alkaline phosphatase or horseradish peroxidase generates chromogenic, target-specific signals that can be analyzed using conventional bright-field microscopes. Additionally, chromatin counterstains were identified to aid in the interpretation of CRISPR-CISH-generated target signals. This advancement makes *in situ* DNA detection techniques more accessible to researchers, diagnostic applications, and educational institutions in resource-limited settings.

## Introduction

Fluorescence *in situ* hybridization (FISH) is a versatile tool for localizing specific nucleic acid sequences in fixed cells and tissues using a fluorescence microscope. Chromogenic *in situ* hybridization (CISH) is an alternative method that detects sequences through chromogenic signal detection, making it compatible with conventional bright-field microscopes (1). The major drawback of standard *in situ* hybridization methods is the need for high temperatures or formamide treatments to denature DNA for probe hybridization, which can damage the chromatin structure (2, 3). Moreover, the hybridization and subsequent wash steps in standard *in situ* hybridization protocols extend the time required to obtain results. To address these issues, a straightforward CRISPR/Cas9-based method called CRISPR-FISH (synonym REGEN-ISL, Ishii et al. 2019) was developed for labeling DNA repeats in fixed plant and animal specimens (4, 5). CRISPR-FISH uses a target-specific crRNA and a universal tracrRNA with a single fluorescent dye added to the 5’ end and dCas9 to label the target sequence (Fig 1A).

**Fig. 1:**
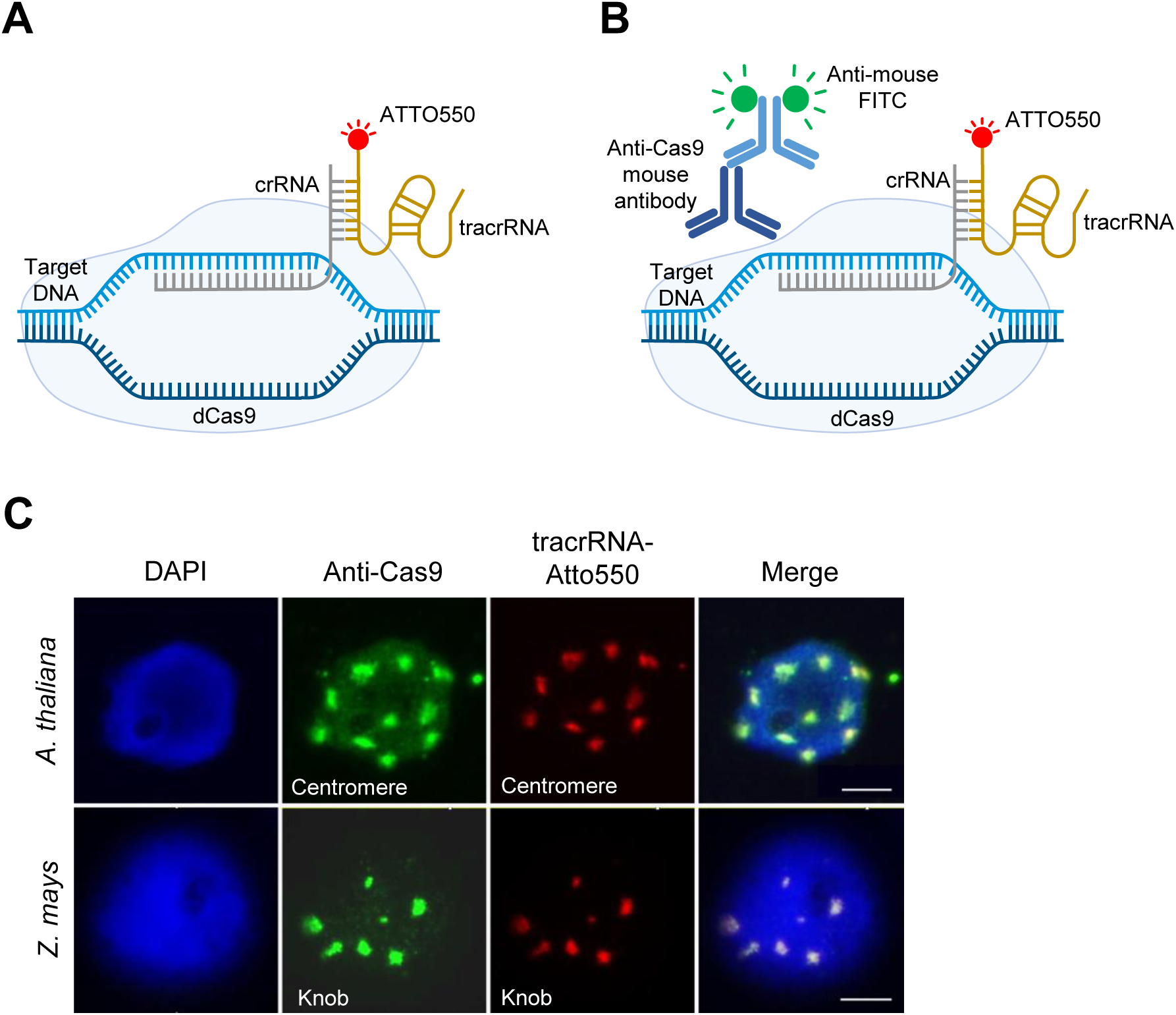
Indirect labeling of DNA repeats by combining CRISPR-FISH with indirect immunostaining. (A) Schematic of standard CRISPR-FISH method. (B) Schematic of indirect CRISPR-FISH to label repeat sequences with Cas9 antibody and later detection with anti-mouse FITC. (C) Visualization of centromere repeats in *A. thaliana* (upper) and knob repeats in *Z. mays* (lower) using CRISPR-FISH and immunofluorescence. Cas9 antibody was detected with anti-mouse FITC (green) and tracrRNA was labeled with Atto550 (red). DNA stained with DAPI (blue) Scale bars, 5 μm.

Unlike most *in situ* hybridization methods, CRISPR-FISH does not require a global DNA denaturation, ensuring a better preservation of the chromatin structure (Němečková et al., 2019). This makes it an excellent tool for studying the three-dimensional genome organization with super-resolution imaging (4–7). The application of the dCas9-RNA complex enables rapid detection of repetitive DNA sequences within seconds, highlighting the fast labeling efficiency of CRISPR-FISH (4). CRISPR-FISH allows multicoloured labeling of repeats in various fixation conditions and specimen samples and works in a large temperature range (4 - 37 °C) (4, 5, 8). Recently, the labeling intensity of CRISPR-FISH has been improved by integrating the ALFA-tag and TSA approaches, making it an even more versatile imaging tool (9). However, broader implementation of this method faces challenges in resource-limited laboratories that lack access to fluorescence microscopes. To overcome this limitation, we developed a robust non-fluorescent CRISPR-labeling method for detecting high-copy repeats. The newly developed CRISPR/Cas9 mediated <u>c</u>hromogenic *in <u>si</u>tu* <u>h</u>ybridization method (CRISPR-CISH) combines chromogenic signal detection methods using alkaline phosphatase or horseradish peroxidase with CRISPR-imaging. The distribution of chromogenic signals can be analyzed by conventional bright-field microscopes. High-copy sequences of mouse and several plant species (*Arabidopsis thaliana, Zea mays*, *Allium fistulosum, Allium cepa* and *Vicia faba*) were used to demonstrate the effectiveness of the developed *in situ* DNA detection method. To assist educators in conducting CRISPR-CISH in their classrooms, a simplified graphical hands-on protocol for schools is provided here.

## Results

### Application of anti-Cas9 for the detection of dCas9-binding sites

High-copy DNA repeats in fixed chromosomes and nuclei can be fluorescently labeled using CRISPR-FISH (4, 5). To explore the possibility of using this CRISPR-based approach to detect repeats non-fluorescently; first, a new CRISPR-FISH strategy had to be developed to label DNA indirectly, as non-fluorescent labeling of DNA can only be achieved indirectly. Therefore, we aimed to test first an indirect CRISPR-FISH method by combining immunofluorescence with standard CRISPR-FISH. In this approach, ATTO550-tagged bipartite gRNA-guided CRISPR-FISH was combined with a Cas9-specific antibody and an anti-mouse conjugated FITC to indirectly visualize dCas9-binding sites (Fig 1B). Using this method, *A. thaliana* centromere and *Z. mays* knob repeats were detected as previously reported using the same crRNAs (Fig. 1C) (4, 7). The overlap of indirect anti-Cas9 (green) and tracrRNA-Atto (red) signals proved the functionality of the tested indirect CRISPR-FISH method (Fig 1C).

### Replacing fluorescence-with non-fluorescence-based labeling of dCas9-binding sites

To replace the fluorescence-based detection of anti-Cas9 binding sites with a non-fluorescence method, we employed alkaline phosphatase (AP) for the detection of the target sequences (Fig 2A). In this approach, anti-mouse conjugated alkaline phosphatase was used to detect anti-Cas9 binding sites. The application of a red chromogenic substrate reacted with alkaline phosphatase, resulting in the expected numbers of red-colored, non-fluorescent signals of centromeres and knob repeat clusters in formaldehyde-fixed nuclei of *A. thaliana* and *Z. mays*, respectively (Fig 2B). Hematoxylin was used as a counter-stain chromatin.

**Fig. 2:**
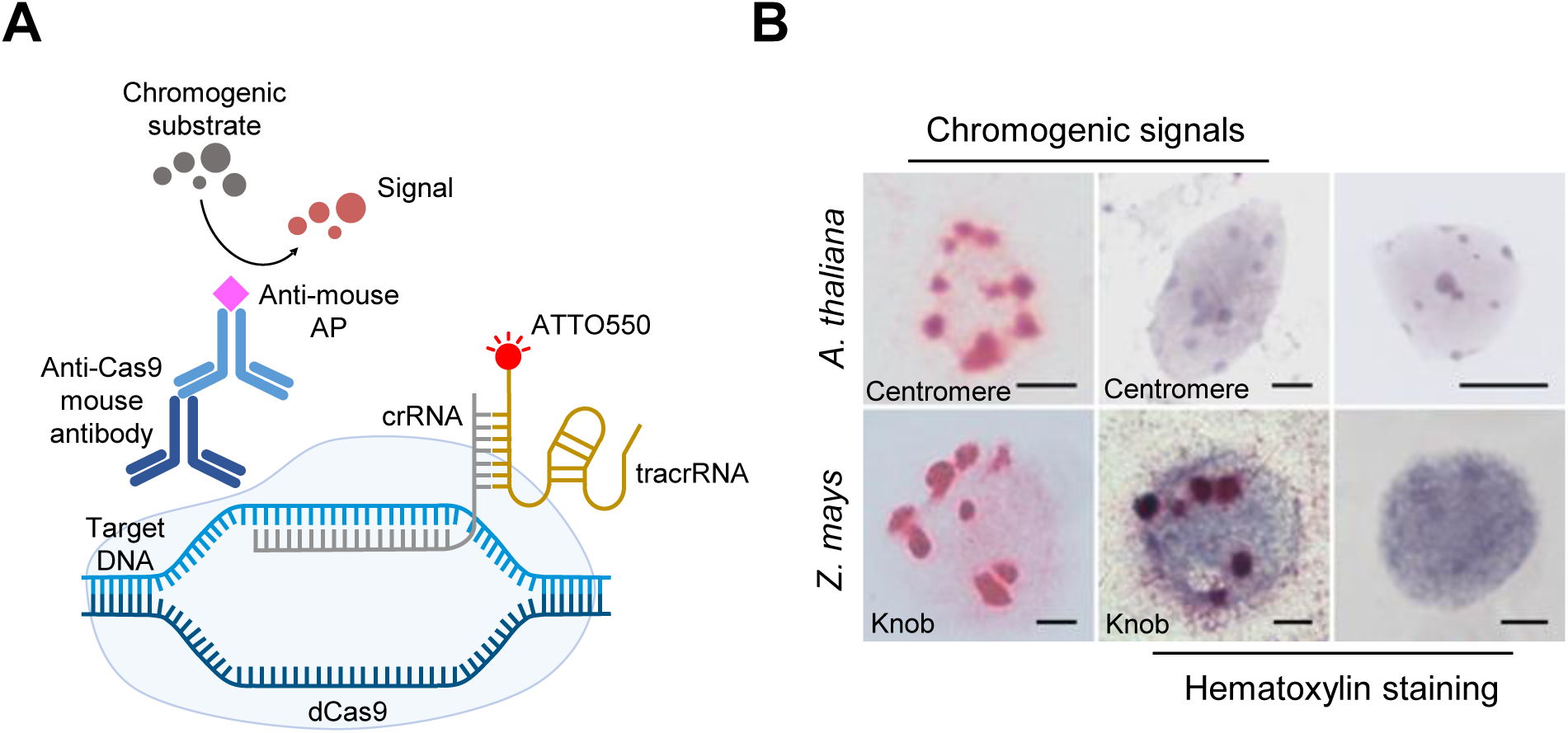
Non-fluorescent labeling of DNA repeats by combining CRISPR-FISH and indirect immunostaining. (A) Schematic of CRISPR-FISH for non-fluorescent labeling of repeat sequences using Cas9 antibody and anti-mouse alkaline phosphatase (AP) and application of red chromogenic substrate develops red colored precipitate at RNP bound sites. (B, upper) non-fluorescent labeling of centromere repeats in *A. thaliana* nuclei (left) without staining, (middle) with hematoxylin counterstaining and (right) with counterstaining only as control without labeling. In *A. thaliana* nuclei, hematoxylin also stained the chromocenters where the actual centromeres were located, so the red chromogenic centromeric dots were masked by hematoxylin. (B, lower) non-fluorescent labeling of knob repeats in *Z. mays* nuclei (left) without staining, (middle) with hematoxylin counterstaining and (right) with counterstaining only as control without labeling. Red-colored spots indicate centromere and knob-specific signals. Scale bars, 5 μm.

However, this dye resulted even in the absence of CRISPR-dCas9 labeling in pronounced staining of the centromeric chromocenters in *A. thaliana* (Fig 2B, upper right), obscuring the detection of the CRIPSR-based centromere signals (Fig 2B, upper middle). Therefore, haematoxylin is not a suitable counterstain for *A. thaliana* nuclei when combined with a non-fluorescent CRISPR-ISH method. In contrast, *Z. mays* nuclei without CRISPR-dCas9 labeling resulted in a more uniform haematoxylin staining (Fig 2B, lower right). The same chromogenic approach without subsequent counterstaining was used to detect *Fok*I and sub-telomeric repeats in *V. faba* and *A. fistulosum* nuclei, respectively (S1A Fig). The observed distinct signal clusters of different sizes were similar to those obtained by CRISPR-FISH control experiments (S1B Fig) with the difference that the non-fluorescent CRISPR-ISH generated a higher background noise level. However, microscopic evaluation of several slides revealed that in all cases, independent of the analyzed species and sequence, only a few nuclei (up to 7.5 %) showed the target-specific signals. This indicates the low efficiency of the indirect detection method using the Cas9 antibody in combination with anti-mouse conjugated alkaline phosphatase. Additionally, these results underscored the need to investigate appropriate nuclear stains across different species to complement non-fluorescent CRISPR-ISH labeling.

### Identification of suitable chromatin counterstains

To enable a correct interpretation of the CRISPR-ISH signals within the analyzed nuclei or chromosomes, we tested chromatin dyes suitable for conventional bright-field microscopy. Out of the six tested dyes, 4% methylene green, 2% methyl green, 4% neutral red, hematoxylin (100% stock solution) and methylene blue (100% stock solution) successfully stained formaldehyde fixed nuclei of *Z. mays*, *V. faba* and *A. fistulosum* within 2 min at RT (S2A Fig). Among these, only neutral red and hematoxylin stained the nuclei in a way that allowed a clear identification of the nuclei in all three species. These two dyes were selected for further experiments. In *A. thaliana,* however, only hematoxylin (100% stock solution) and methylene blue (100% stock solution) effectively stained the nuclei within 10 min of dye application (S2B Fig). Hematoxylin provided stronger staining compared to methylene blue, but both dyes intensely stained chromocenters (S2B Fig), rendering them unsuitable as they could obscure CRISPR-ISH signals. Thus, we decided not to use a chromatin dye in *A. thaliana* for the subsequent experiments. In general, to achieve a clear contrast between CRISPR-ISH signals and the chromatin stain, it is essential to select an appropriate counterstain based on the chromogenic substrate used.

Moreover, our observations revealed a likely correlation between the counterstaining ability of chromatin and the genome size of the species. For instance, the small-genome species *A. thaliana* (142.5 Mb/1C; https://www.ncbi.nlm.nih.gov/datasets/genome/GCA_028009825.2/) exhibited weaker chromatin staining compared to the large-genome species *V. faba* (13 Gb/1C) (10) and *A. fistulosum* (11.27 Gb/1C) (11).

### The application of biotinylated tracrRNA improved indirect CRISPR-ISH and enabled CRISPR-CISH

To increase the efficiency of the indirect CRISPR-ISH approach, we first employed 3’ biotin-labeled tracrRNA in combination with streptavidin-conjugated FITC (Fig 3A). To assay the functionality of this approach, *Arabidopsis* centromere-and *Arabidopsis*-type telomere-specific crRNAs were applied and formaldehyde fixed *A. thaliana* and *N. benthamiana* nuclei showed target sequence-specific labeling, respectively (Fig 3B). As a positive control, standard CRISPR-FISH using Atto550-labeled tracrRNA and standard FISH with biotin-labeled oligo-probes specific for both sequences were used (S3 A & B Fig). In addition, the specificity of the indirect CRISPR-FISH signals was demonstrated by probing *A. thaliana* centromere and *Z. mays* knob repeats by simultaneous application of the standard CRISPR-FISH and indirect CRISPR-FISH (Fig 3C). To compare the labeling efficiency of both methods, centromere and knob signals of 2C sorted nuclei generated by both standard and indirect CRISPR-FISH were quantified. In the counting assay, about 8 and 7 centromere-specific signals in *A. thaliana* (Fig 3D, left) and about 6 and 5 knob-specific signals in *Z. mays* (Fig. 3D, right) were detected by standard and indirect CRISPR-FISH, respectively. In addition, 47 and 45 nuclei out of 50 *A. thaliana* nuclei showed centromere-specific signals and 45 and 43 nuclei out of 50 *Z. mays* nuclei showed knob specific-signals generated by standard and indirect CRISPR-FISH, respectively (S3C Fig). Overall, the quantification results showed a satisfactory labeling efficiency of indirect CRISPR-FISH using a 3’ biotin-labeled tracrRNA compared with standard CRISPR-FISH.

**Fig. 3:**
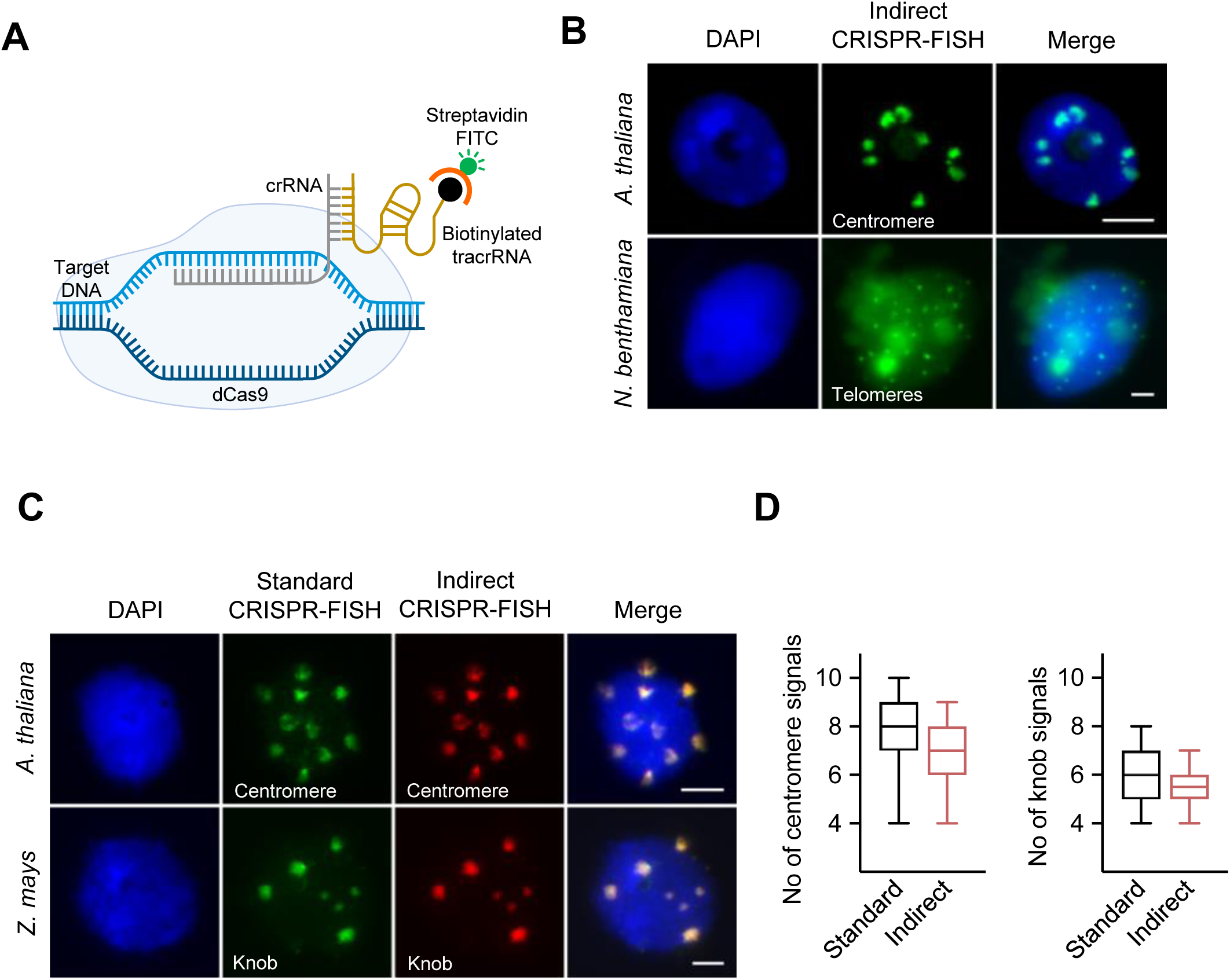
Indirect CRISPR-FISH labeling of DNA repeats using biotinylated tracrRNA. (A) Schematic of indirect CRISPR-FISH for the labeling of repeat sequences using biotinylated tracrRNA and detection with streptavidin FITC. (B) Visualization of (upper) centromere repeats in *A. thaliana* and (lower) telomere repeats in *N. benthamiana* (bottom) with indirect CRISPR-FISH using biotinylated tracrRNA and detected with streptavidin FITC. (C) Images showing colocalization of indirect CRISPR-FISH (green) and standard CRISPR-FISH (red) spots labeling centromeres and knob repeats in *A. thaliana* and *Z. mays* nuclei, respectively. Biotinylated tracrRNA was detected using streptavidin FITC (green). (D) Comparison of labeling efficiency between two methods, number of (left) centromere and (right) knob signals generated per nucleus by standard CRISPR-FISH and indirect CRISPR-FISH methods, n =50 nuclei. Upper and lower whiskers indicate the maximum and minimum number of signals observed. The black line in the middle of each box plot indicates the median. DNA stained with DAPI (blue). Scale bars, 5 μm.

Finally, we employed the 3’ biotin-labeled tracrRNA in combination (I) with streptavidin-conjugated alkaline phosphatase (Fig 4A) or (II) the streptavidin-conjugated horseradish peroxidase (Fig 4C) and corresponding chromogenic substrates. We termed this strategy “CRISPR-CISH”, a <u>CRISPR</u>-Cas9 mediated <u>c</u>hromogenic *in <u>s</u>itu* <u>h</u>ybridization method. The application of the streptavidin-conjugated alkaline phosphatase enzyme and the red substrate for signal detection in formaldehyde-fixed *A. thaliana* and *Z. mays* nuclei using centromere and knob-specific gRNA, respectively, resulted in a specific labeling of the repeats (Fig 4B). Although the signals generated by CRISPR-CISH in combination with alkaline phosphatase were specific, the structure of the signals was a bit fuzzy. In contrast, the application of the peroxidase enzyme along with the green chromogenic substrate resulted in clear and sharper green-colored centromere and knob signals (Fig 4D). After the chromogenic reaction, *Z. mays* nuclei were counterstained in 4% neutral red for 2 mins. Using a conventional bright-field microscope revealed strong green knob-specific signals on light orange-stained nuclei (Fig 4E, upper), as a control *Z. mays* nucleus were counterstained without CRISPR-CISH (Fig. 4E, lower). To assess the efficiency of CRISPR-CISH, we compared the number of knob repeat-specific signals generated by indirect CRISPR-FISH and CRISPR-CISH. About 5 knob-specific signals were counted after indirect CRISPR-FISH and CRISPR-CISH, respectively (Fig 4F). In addition, 43 and 41 nuclei out of 50 *Z. mays* nuclei showed knob specific-signals generated by indirect CRISPR-FISH and CRISPR-CISH, respectively (Fig 4G).

**Fig. 4:**
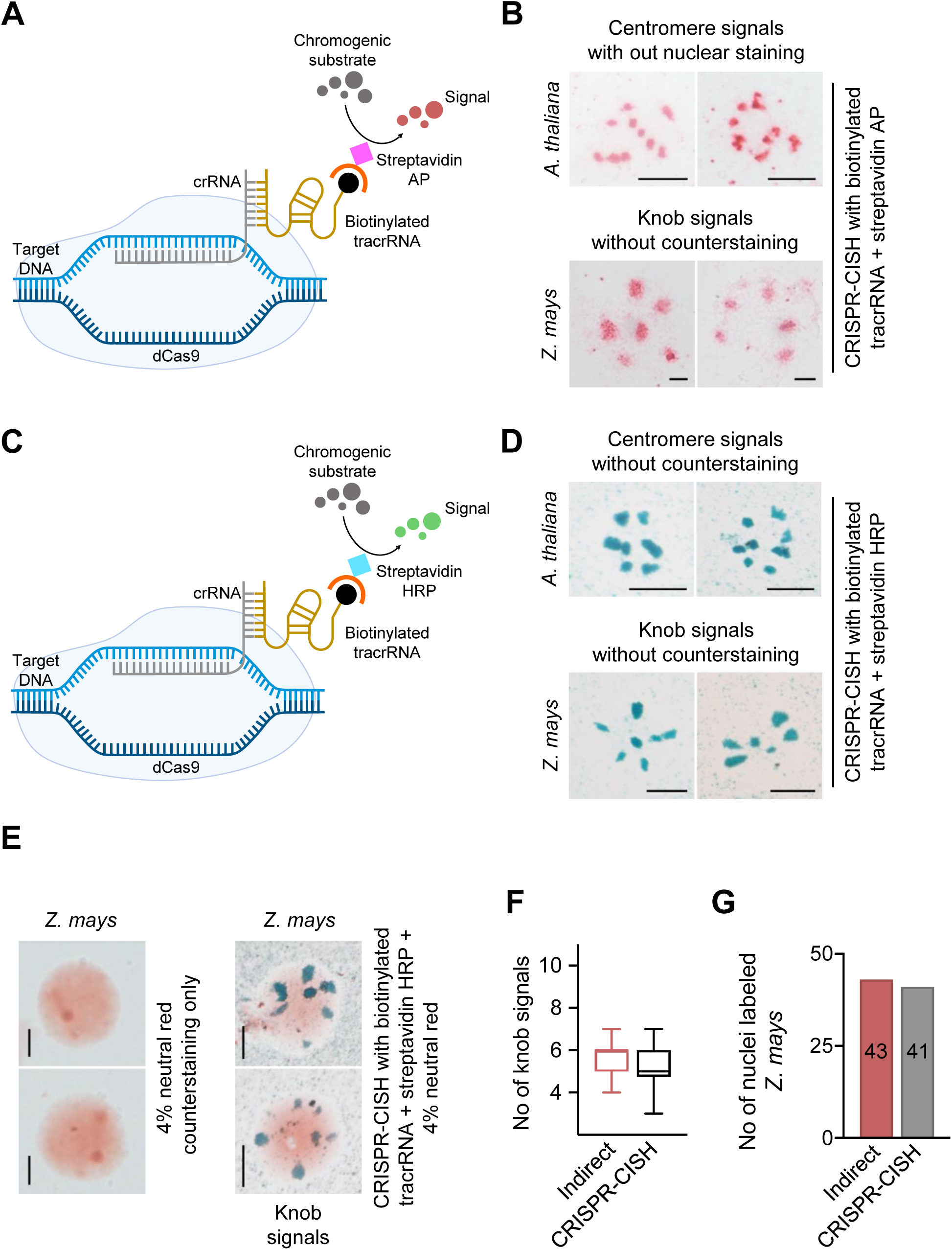
CRISPR-Cas9-mediated chromogenic *in situ* detection of repeat sequences in fixed samples. (A) Schematic of CRISPR-CISH for labeling repeat sequences using biotinylated tracrRNA and streptavidin alkaline phosphatase (AP) and application of red chromogenic substrate develops red colored precipitate at RNP bound sites. (B) CRISPR-CISH-based non-fluorescent labeling of centromeres (top) and knob repeats (bottom) in *A. thaliana* and *Z. mays* nuclei without counterstaining using streptavidin AP and red chromogenic substrate. Red-colored spots indicate centromere and knob-specific signals. (C) Schematic of CRISPR-CISH for labeling repeat sequences using biotinylated tracrRNA and streptavidin-horseradish peroxidase (HRP) and application of green chromogenic substrate develops green colored precipitate at RNP bound sites. (D) CRISPR-CISH-based non-fluorescent labeling of centromeres (top) and knob repeats (bottom) in *A. thaliana* and *Z. mays* nuclei without counterstaining using streptavidin HRP and green chromogenic substrate. Green-colored spots indicate centromere and knob-specific signals, respectively. (E, upper) CRISPR-CISH labeling of knob repeats in *Z. mays* nuclei counterstained with 4% neutral red and (E, lower) counterstained only as a control with no labeling. (F) Comparison of labeling efficiency between two methods, number of knob signals generated per nucleus by indirect CRISPR-FISH and CRISPR-CISH methods, n =50 nuclei. Upper and lower whiskers indicate the maximum and minimum number of signals observed. The black line in the middle of each box plot indicates the median. Scale bars, 5 μm. (G) Bar graph comparing the efficiency of indirect CRISPR-FISH and CRISPR-CISH methods. The number of nuclei with knob repeats on fixed 2C nuclei of *Z. mays* is shown. n =50 nuclei.

Finally, to test whether also CRISPR-CISH labels repeats on conventionally 3:1 (ethanol: acetic acid) fixed chromosomes and nuclei, we employed *Z. mays* and *V. faba* chromosomes for labeling the knob and *Fok*I repeats, respectively. After CRISPR-CISH application, either 4% neutral red or 100% hematoxylin was used to stain the chromosomes. CRISPR-CISH successfully labeled knob repeats in *Z. mays* chromosomes, and specific signals were observed at the terminal regions of the chromosome arms (Fig 5A, left). Also, CRISPR-CISH labeled *Fok*l repeats in the mid-arm position of the long arms of all *V. faba* chromosomes (Fig 5A, right). The application of 4% neutral red after CRISPR-CISH resulted in both species in light red counterstained chromosomes. The application of hematoxylin in combination with CRISPR-CISH exhibited green counterstained chromosomes in both species (Fig 5B). Using CRISPR-CISH in combination hematoxylin also successfully labeled the centromere repeats of *A. thaliana,* sub-telomeric repeats of *A. fistulosum* and the major satellite repeats of mouse chromosomes (Fig 5C). As a positive control, streptavidin-conjugated FITC was used to detect indirect CRISPR-FISH specific signals in 3:1 fixed *A. thaliana*, *Z. mays*, *V. faba*, *A. fistulosum*, and mouse chromosomes (S4 Fig).

**Fig. 5.**
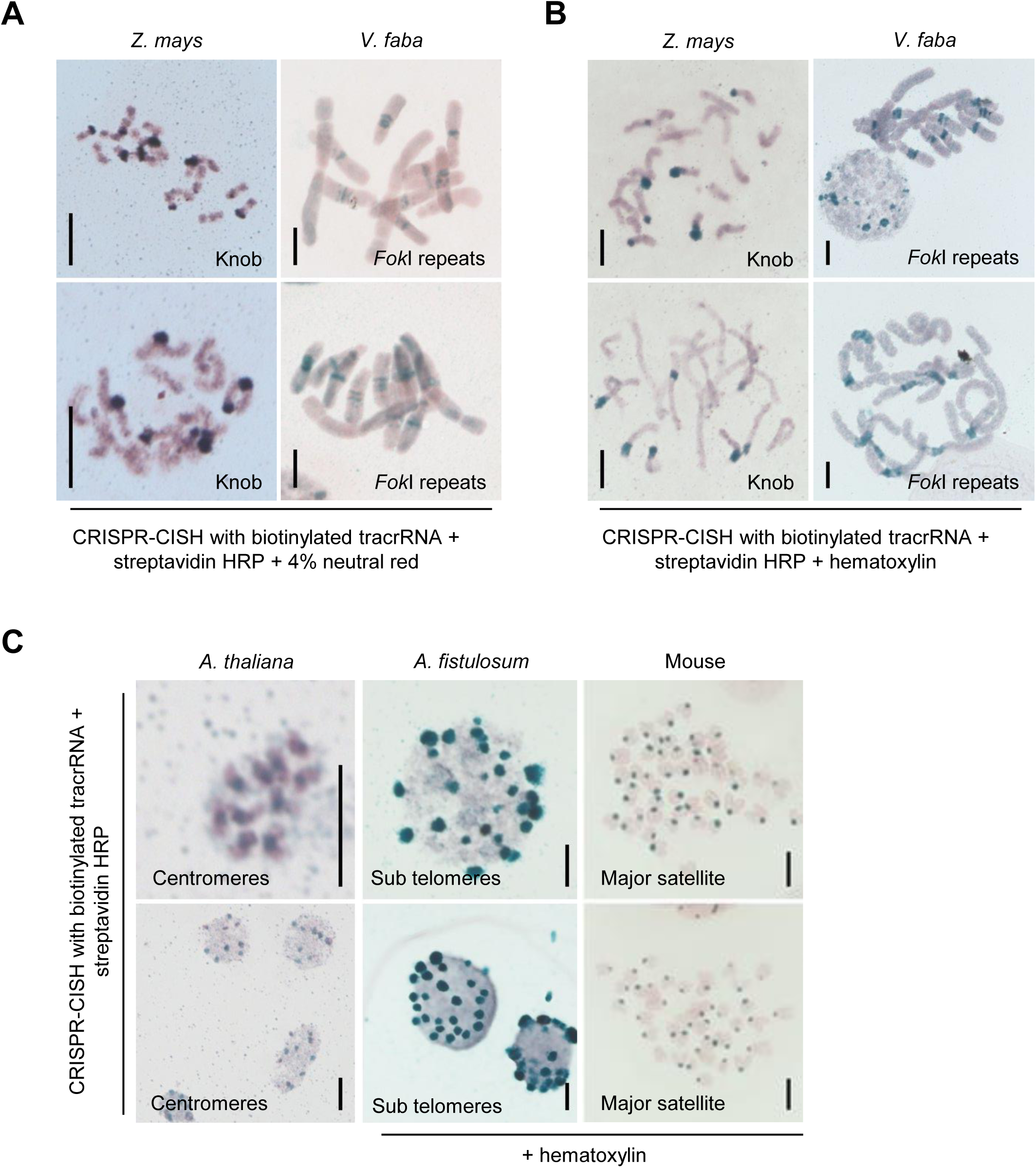
CRISPR-Cas9-mediated chromogenic *in situ* detection of repetitive sequences in ethanol: acetic acid fixed chromosomes and nuclei. Non-fluorescent visualization of knob and *Fok*I repeats on *Z. mays* (left) and *V. faba* (right) chromosomes using the CRISPR-CISH method in combination with (A) neutral red and (B) hematoxylin staining. (C) Visualization of centromere, sub telomere and major satellite repeats in *A. thaliana*, *A. fistulosum* and mouse on ethanol: acetic acid fixed chromosomes using CRISPR-CISH. *A. fistulosum* and mouse samples were stained with hematoxylin. Scale bars, 5 μm.

In conclusion, CRISPR-CISH enables the successful labeling of repeats in both formaldehyde or 3:1 fixed nuclei and chromosomes. Conventional bright-field microscopy is sufficient for the analysis of the specimens. The application of horseradish peroxidase-based detection is better than alkaline phosphatase because it results in more distinct and clear chromogenic signals with less background. A detailed protocol for the application of CRISPR-CISH in educational institutes and science outreach settings is provided in www.protocols.io, DOI: dx.doi.org/10.17504/protocols.io.kqdg3q1×1v25/v1 (www.protocols.io/private/D61DAFC0E91011EFBAE60A58A9FEAC02).

## Discussion

Standard CRISPR-FISH uses a fluorescently labeled tracrRNA to visualize repetitive DNA sequences in fixed samples (4). In our study, we developed a non-fluorescent CRISPR-ISH method and evaluated two indirect labeling strategies: using anti-Cas9 antibodies and biotinylated tracrRNA. The Cas9 antibody approach was less efficient, likely due to formaldehyde fixation, which may alter protein conformations and mask epitopes, hindering the accessibility of the antibody (12, 13). In contrast, the application of biotinylated tracrRNA resulted in a higher labeling efficiency. The strong and specific interaction between biotin and streptavidin remained stable even under stringent fixation and washing conditions, contributing to the enhanced performance of this method (14). When using biotinylated tracrRNA, CRISPR-CISH produced much clearer signals with the streptavidin-horseradish peroxidase conjugate than with the alkaline phosphatase conjugate. This could be due to the larger size of the alkaline phosphatase (approximately 86 kDa), which may hinder its ability to penetrate the chromatin (15). In contrast, horseradish peroxidase, which is only 40 kDa in size, likely penetrates the chromatin more efficiently (16).

Compared to other tested chromatin counterstains, haematoxylin and neutral red showed optimal staining, providing better contrast between signals and nuclei or chromosomes. Unfortunately, none of the non-fluorescence dyes stained *A. thaliana* nuclei uniformly, indicating the need for further investigations by screening additional dyes in future studies.

Unlike standard FISH, CRISPR-CISH is a fast and convenient method that offers several advantages over conventional DNA detection techniques. In particular, the application time of CRISPR-CISH is much shorter due to the rapid DNA binding of Cas9 (4). In contrast to standard FISH, which requires an expensive fluorescence microscope, CRISPR-CISH only needs a conventional light microscope to visualize the labeled DNA. In addition, the chromogenic signals generated by CRISPR-CISH are sufficiently strong to be visualized with a 40x lens without fading over time, allowing specimens to be archived for long periods. Moreover, CRISPR-CISH uses mild conditions that make it easy and safe to conduct hands-on investigations in educational institutions or laboratories where expensive fluorescence microscopes and specialists may not be available.

In educational settings, CRISPR-CISH can be a powerful tool to inspire students by enabling them to visualize DNA repeats under a conventional bright-field microscope. To make CRISPR-CISH more accessible in schools, it could be combined with the Foldscope, a paper-based origami microscope that offers 140x to 350x magnification and is available as a classroom kit (https://foldscope.com) (17). Additionally, using pre-assembled dCas9/gRNA RNP complexes to label DNA eliminates the need for GMO-related permissions and raises no ethical concerns, making it an ideal tool for classroom applications.

However, the application of CRISPR/Cas9 is limited by the requirement for the NGG PAM sequence at the target site, which is necessary for designing the gRNA. This limitation can be overcome by using engineered Cas9 variants that work with different PAM sequences (reviewed by (18)). To date, CRISPR-CISH has efficiently labeled high-copy genomic DNA repeats, but it is less efficient at labeling low- and single-copy sequences. To improve its utility for low-copy sequences, CRISPR-CISH could potentially be combined with the dCas9-ALFA-tag approach, which has recently demonstrated increased signal intensities (9). Despite these limitations, CRISPR-CISH holds significant potential for future research and diagnostic applications in less-funded settings, where it can facilitate the investigation of basic chromosome biology.

## Materials and Methods

Nuclei and chromosomes from *Arabidopsis thaliana*, maize (*Zea mays*), Welsh onion (*Allium fistulosum*), onion (*Allium cepa*), broad bean (*Vicia faba*), and the house mouse (*Mus musculus forma domestica*) were used.

### Preparation of leaf nuclei

Young leaf tissue was fixed in formaldehyde (2% or 4%) in ice-cold Tris buffer (10 mM Tris-HCl, 10 mM Na_2_-EDTA, 100 mM NaCl, 0.1% Triton X-100, and adjusted pH 7.5 with NaOH) under vacuum for 5 min and followed by incubation for 25 min on ice without vacuum. The tissue was rinsed twice in ice-cold Tris buffer for 5 min each. Then, the tissue was chopped in a Petri dish in 300 - 500 µl ice-cold LB01 buffer (15 mM Tris-HCl (pH 7.5), 2 mM Na_2_-EDTA, 0.5 mM spermin, 80 mM KCl, 20 mM NaCl, 15 mM β-mercaptoethanol and 0.1% Triton X-100) using a fresh razor blade. The suspension was filtered through a 35 µm cell-strainer cap (BD Biosciences) and spun onto glass slides using a Cytospin 3 (Shandon) with 700 rpm for 5 min. The slides were either used immediately or kept in 1x PBS at 4°C overnight. Alternatively, slide preparation with formaldehyde-fixed cells could be performed using a sucrose buffer instead of a cytospin, as described by (19).

### Preparation of chromosomes

Root tip meristems were obtained from 3 - 5 days old *Z. mays, V. faba* and *A. cepa* seedlings and potted *A. fistulosum* plants. *Z. mays and V. faba* roots were treated in 0.1% colchicine at RT for 3 h. *A. cepa* and *A. fistulosum* roots were treated in ice-cold water for 24 h. Then, roots of all species were fixed in 3:1 (ethanol: acetic acid) solution for 24 h at RT and washed twice in ice-cold ddH_2_0 and 0.01 M citrate buffer (0.01 M sodium citrate and 0.01 M citric acid, pH 4.5–4.8), for 5 min each. 3 - 5 root tips of *Z*. *mays, V. faba, and A. fistulosum* were digested for 50, 60 and 60 min, respectively in 25 – 50 µl of enzyme mixture depending on the number of root tips (0.7% cellulase R10, 0.7% cellulase, 1% pectolyase, 1% cytohelicase dissolved in 0.01 M citrate buffer) at 37 ⁰C. The tubes with digested meristems were vortexed and, after adding 600 µl of ddH_2_O centrifuged at 10,000 rpm for 45 s. The supernatant was removed, and 600 µl 96% ethanol was added and centrifuged at 11,000 rpm for 30 s. After the supernatant was discarded by inverting the tubes, the pellets were resuspended in 80 – 100 µl 96% ethanol. For the preparation of chromosome slides, 10 µl of cell suspension was dropped from a height of 10 – 15 cm onto the glass slide in a moisture box placed on a hot plate at 55° C. Then 25 – 50 µl of freshly prepared ethanol: acetic acid (3:1) at room temperature (RT) was added to the glass slide and air dried.

*A. cepa* chromosomes were prepared using the squash technique. A 3:1 fixed single root was incubated in 45% acetocarmine in acetic acid for 5 minutes. The root cap was removed, and the meristematic tip was squeezed onto a slide. After adding 10 - 15 µl of 45% acetic acid, the sample was covered with a coverslip and gently tapped. Brief heating was applied with a spirit burner from beneath the slide, followed by freezing with liquid nitrogen. The coverslip was then removed, and the slide was immediately immersed in 99% ethanol and then air-dried.

*A. thaliana* chromosomes were obtained from young flower buds. The flower buds were fixed in 3:1 (ethanol: acetic acid) solution for 24 h at RT. Then, the flower buds were washed twice in ddH_2_0 and buds containing yellow anthers were removed. The remaining buds were washed twice in 0.01 M citrate buffer for 5 min each on ice and digested with a 50% enzyme mixture (0.35% cellulase R10, 0.35% cellulase, 0.5% pectolyase, 0.5% cytohelicase dissolved in 0.01 M citrate buffer) in 1x citrate buffer at 37 ⁰C for 75 min. Later, the enzyme mixture was replaced with 0.01 M citrate buffer. For the preparation of chromosome slides, a single flower bud was placed onto a glass slide and 10 - 20 µl of 60% acetic acid was added. The slide was placed on a hot plate (50 ⁰C) and the suspension was spread by circular stirring with a dissecting needle for 30 s (20). Later, 25 - 50 µl of freshly prepared ethanol: acetic acid (3:1) was added to the glass slide and dried. Slides containing well-spread chromosomes were selected using a phase-contrast light microscope.

Murine chromosomes were obtained from the skin of a laboratory house mouse strain C57Bl6/J. Chromosome slide preparation was done as described in (5).

### Preparation of recombinant dCas9

The *Streptococcus pyogenes* (dead version) Cas9 gene sequence containing the double nuclease mutation (D10A and H840A) was PCR-amplified from the dCas9:3×PP7:GFP vector (21) and cloned into pET22b+ (Invitrogen) with a C-terminal hexahistidine affinity tag. The plasmid was transformed into *Escherichia coli* BL21 (DE3) and grown in 2x TY media at 37°C to OD600 ∼ 0.5 and then induced by 0.5 mM isopropyl-β-D-1-thiogalactopyranoside (IPTG) for 16 h at 18°C. Cells were lysed in lysis buffer (50 mM NaH_2_PO_4_, 500 mM NaCl, 10% glycerol, 10 mM imidazole, pH 8) containing lysozyme (1 mg/ml) for 30 minutes on ice, then snap frozen in liquid nitrogen and immediately thawed in water at room temperature (RT) followed by sonication on ice for 4 times for 30 seconds each at 50% intensity using a Vibra-Cell sonicater Model VC60 (Sonics & Materials Inc.). The lysate was centrifuged at 7000 rpm for 20 min at 4°C, and the supernatant was transferred to a Falcon tube containing 1 ml of PureCube 100 Ni-NTA Agarose (Cube Biotech, 31103) and rotated at 4°C for 90 min. The His-tagged dCas9 protein was then purified by gravity flow chromatography by passing the supernatant through disposable polypropylene columns (Qiagen, 34924) and eluted with elution buffer (50 mM NaH_2_PO_4_, 500 mM NaCl, 10% glycerol, 250 mM imidazole, pH 8). Purified dCas9 protein was stored at -20°C. Alternatively, recombinant dCas9 can be commercially purchased, e.g. from Novateinbio (Catalog: PR-137213).

### Standard CRISPR-FISH

We employed the bipartite gRNA (crRNA and 3’ATTO 550/Alexa Fluor 488 –labeled tracrRNA) system (Alt-R® CRISPR-Cas9, Integrated DNA Technologies, https://eu.idtdna.com) for CRISPR-FISH (synonym REGEN-ISL, Ishii et al., 2019). Target-specific crRNA were designed using the software CRISPRdirect (https://crispr.dbcls.jp/) (S1 Table). Preparation of the RNP complex was done using the standard protocol (4). Prepared slides were washed with 0.2% Triton-X100 in Tris-HCL (pH 9) at 37 °C for 1 – 2 h, and later washed in 1x PBS twice for 5 min each. 100 µl of 1x Cas9/1mM DTT buffer (200 mM Hepes (pH 7.5), 1M KCl, 50 mM MgCl_2_, 50% glycerol, 10% BSA, and 1% Tween 20) was added per slide for 5 min at RT and then buffer was removed by tilting the slides. 20 - 25 µl of RNP complex was added per slide and covered with parafilm and incubated at 37 °C for 1 hour in a moisture box. After incubation, the slides were washed in 1x PBS for 5 min and post-fixed with 4% formaldehyde in 1x PBS for 5 min at RT. Then the slides were washed with 1x PBS and dehydrated in an ethanol series (70%, 90%, and 96%) for 2 min each. Finally, slides were air-dried in the dark, and counterstained with 8 µl of (0.5 μg/ ml) 4′,6-diamidino-2-phenylindole (DAPI) in antifade (Vector Laboratories, Burlingame, CA, USA) and stored at 4 °C for further microscopy.

### Indirect CRISPR-FISH employing an anti-Cas9 antibody

After standard CRISPR-FISH, slides were blocked with 100 µl of 4% BSA for 1 h at RT in the dark. Then 50 µl of a monoclonal anti-Cas9 mouse antibody (Santa Cruz Biotechnology, 7A9-3A3) diluted 1:300 in 2% BSA in 1x PBS was added per slide and incubated at 37 °C for 1 h and subsequently overnight at 4 °C in a humid box. The slides were washed in 1x PBS twice for 5 min on ice. Then 50 µl of secondary anti-mouse Alexa 488 antibody (Thermo Fisher Scientific Inc. Massachusetts, U.S.A. cat: A11001) diluted 1:200 in 2 % BSA in 1x PBS was applied per slide and incubated at 37 °C for 1 h. Subsequently, the slides were washed twice with ice-cold 1 x PBS for 5 min. Dehydration and counterstaining were performed as described for standard CRISPR-FISH.

### Indirect CRISPR-FISH using biotinylated tracrRNA

A 3’ biotinylated tracrRNA (Integrated DNA Technologies, https://eu.idtdna.com, S1 Table) and target-specific crRNA were used for the preparation of the RNP complex using a standard protocol (4). CRISPR-FISH was performed as described for standard CRISPR-FISH until post-fixation and washing for 5 min in 1x PBS at RT. Afterwards, the slides were blocked with 100 µl of 4% BSA in 1x PBS for 30 min at RT and then washed twice in 1x PBS for 5 min each. Then 50 µl of streptavidin-conjugated FITC (Sigma-Aldrich, S3762) diluted 1:40 in 2% BSA/1x PBS was added per slide and incubated at 37 °C for 1 h. Subsequently, the slides were washed twice in 1 x PBS for 5 min each at RT in dark and dehydrated and counterstained as described for standard CRISPR-FISH.

### CRISPR-CISH - CRISPR Cas9 mediated chromogenic *in situ* **detection**

CRISPR-CISH was performed like indirect CRISPR-FISH using biotinylated tracrRNA till post-fixation and washing for 5 min in 1x PBS at RT. The slides were blocked with 100 µl of 4% BSA in 1x PBS for 30 min at RT and then washed twice in 1x PBS for 5 min each. Then 50 µl of streptavidin-conjugated alkaline phosphatase (AP) (ZytoChem Plus AP Kit, AP008RED) or horseradish peroxidases (HRP) (Permanent HRP Green Kit, ZUC070-100) per slide was added according to manufacturer’s instructions and incubated at 37 °C for 1 h. Then, the slides were washed twice in 1x PBS for 5 min each at RT. In the case of AP, 50 µl of permanent red buffer with 0.8 µl permanent red concentrate (ZytoChem Plus AP Kit, AP008RED) was added and incubated at RT until the red color developed (usually around 10 minutes, the precise time should be figured out empirically). In the case of HRP, 50 µl of HRP green substrate buffer with 4.5 µl HRP green chromogen (Permanent HRP Green Kit, ZUC070-100) was added and incubated at RT until the green color developed (over 10 minutes). The desired color intensity was monitored using a light microscope. Then, the slides were washed in ddH_2_O at sufficient color intensity. The slides were counterstained with a nuclear dye and later rinsed with tap water and air-dried. Finally, slides were mounted with Entellan® (Merck KGaA) and analyzed by light microscopy.

### Standard fluorescence *in situ* hybridization (FISH)

FISH with a 5′ biotin-labeled *A. thaliana* centromere-specific (22) and telomere-specific oligo probes (23) (S1 Table) was performed as described in (24). Nuclei-carrying slides were prepared as mentioned for CRISPR-FISH. Then, they were washed with 2x SSC for 5 min each and further treated with 45% acetic acid for 10 min at RT. Afterwards, slides were washed in 2x SSC for 5 min each and fixed in 4% formaldehyde in 1x PBS for 10 min at RT. To remove the excess of fixative, the slides were washed thrice with 2x SSC for 4 minutes each and dehydrated in an ethanol series (70%, 90%, and 96%) for 2 min each, followed by air drying. 20 μl of hybridized buffer (50% deionized formamide, 20x SSC, 10% dextran) containing probe was added on each slide under a 22 x 22 coverslip and denatured at 80 °C for 2 min on a hot plate and incubated overnight at 37 °C in a moisture box. On the next day, the slides were washed twice in 2x SSC for 5 min each to remove the coverslip. To remove the unspecifically bound probes, slides were washed with 2x SSC at 58 °C for 20 min in a water bath and later transferred to 2x SSC at RT for 2 min. Then the slides were incubated with 100 μl of blocking solution (4% BSA, 0.1% Tween 100, 1x PBS) under parafilm for 30 at RT and washed twice for 5 min each in 2x SSC at RT. Afterwards, slides were probed with streptavidin conjugated to FITC (Sigma-Aldrich) (1:100 diluted) for 1 h at 37°C under parafilm in a humid chamber, followed by washing thrice for 5 min each in 2x SSC at RT in the dark and dehydrating in an ethanol series (70%, 90%, and 96%) for 2 min each. Finally, the slides were air-dried in the dark and counterstained with 8 µl of (0.5 μg / ml) 4′,6-diamidino-2-phenylindole (DAPI) in antifade (Vector Laboratories, Burlingame, CA, USA) and stored at 4 °C for further microscopy.

### Counterstaining of specimens with nuclear dyes

Methylene green, methyl green (Sigma, M8884-5G), neutral red (Feinchemie K.-H. Kallies KG), and naphthol green B (Waldeck, 1B-385) solutions were prepared in ddH_2_O. In addition, 100% stock solutions of hematoxylin (Sigma, GHS3-50ML), and methylene blue (Merck, 115943) were tested. 300 µl of dye was added to each slide for counterstaining.

### Microscopy

Chromogenic *in situ* imaging was performed using an Axiophot light microscope (Carl Zeiss) equipped with an Axiocam 506 color camera and operated using Zen 3.0 blue Edition software (Carl Zeiss AG. Germany). Fluorescence imaging was performed using a BX61 microscope (Olympus) equipped with an ORCA-R2 CCD camera (Hamamatsu) and operated using cellSens imaging software. All images were captured in grayscale and later pseudo-colored using ImageJ software (https://imagej.nih.gov/ij/).

### Signal quantification

For quantification of signals, leaf tissues of *A. thaliana* and *Z. mays* were fixed in 4% and 2% formaldehyde in TRIS buffer, respectively (4, 7), respectively. Nuclei were isolated and 2C nuclei were sorted using a BD Influx cell sorter (BD Biosciences) according to (25). CRISPR-imaging signals from 50 flow-sorted 2C nuclei per each analysis were counted.

## Acknowledgements

We would like to thank Hans Peter Mock, former principal investigator of the research group “Applied Biochemistry” (IPK Gatersleben, Germany), for his help in protein purification. We also thank Thomas Liehr (Friedrich Schiller University, Jena) for providing us with mouse chromosome suspension and Petra Gerstmann for sharing hematoxylin.

## Author contributions

AH and BPP designed the research. BPP performed the majority of the experiments. FD performed indirect CRISPR labeling experiments using a Cas9 antibody. DRS tested different nuclear stains. JF performed flow sorting. BPP, JF, and AH wrote the manuscript.

## Data availability

All data supporting the findings of this study are available within the paper and within its supplementary materials published online. Methods are described in in www.protocols.io, DOI: dx.doi.org/10.17504/protocols.io.kqdg3q1×1v25/v1 (www.protocols.io/private/D61DAFC0E91011EFBAE60A58A9FEAC02).

## Supporting information

**S1 Fig:**
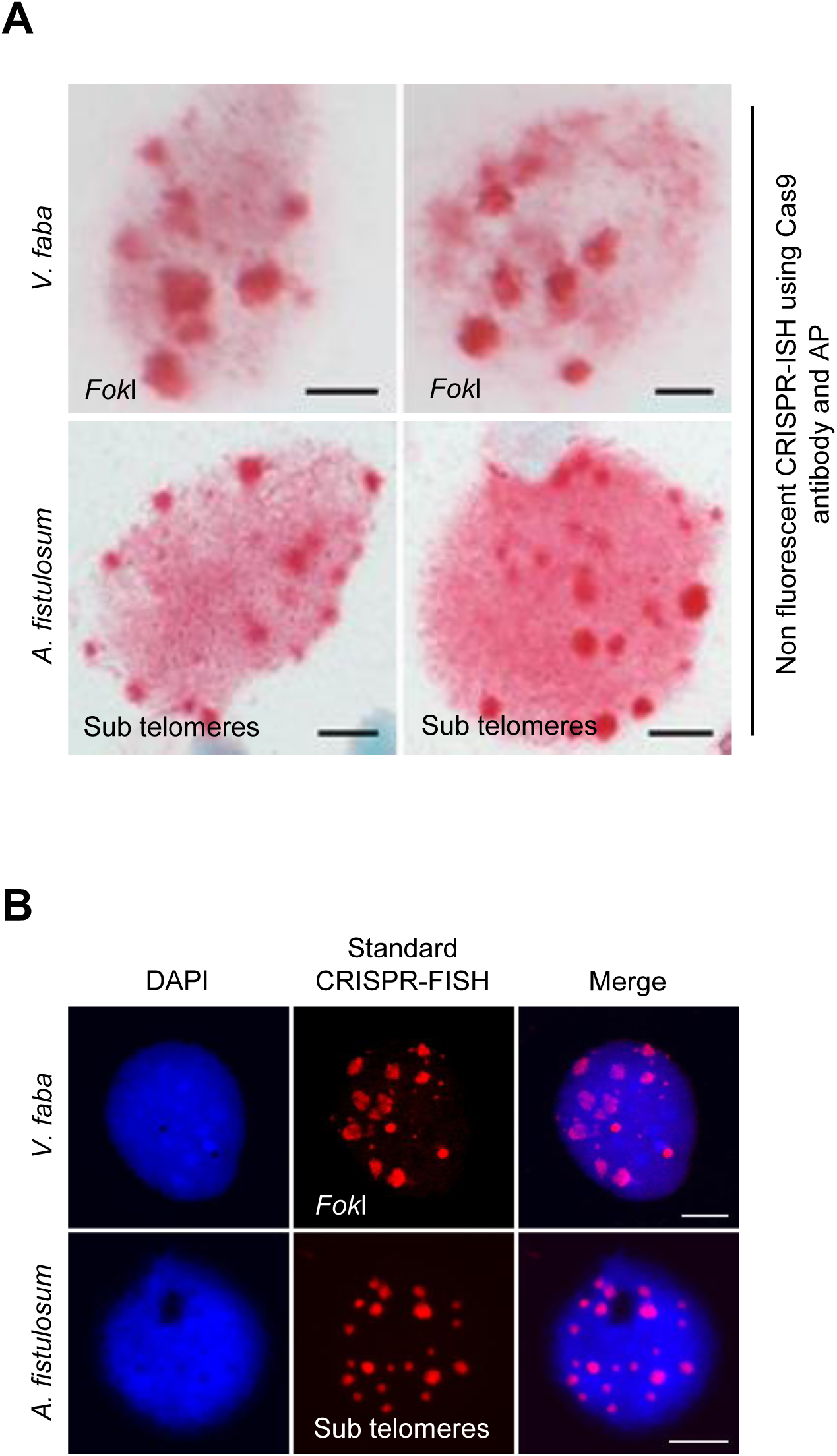
(A) Non-fluorescent labeling of DNA repeats by combining CRISPR-ISH with immunoassay on formaldehyde fixed nuclei of *V*. *faba* (upper) and *A. fistulosum* (lower), labeling *Fok*I and sub telomeres, respectively, without counterstaining. Dark red spots indicate *Fok*I and sub-telomeric specific signals, respectively. (B) Control labeling of FokI and sub-telomeric repeats on formaldehyde-fixed nuclei of *V. faba* (upper) and *A. fistulosum* (lower) using CRISPR-FISH. Scale bars, 5 μm.

**S2 Fig:**
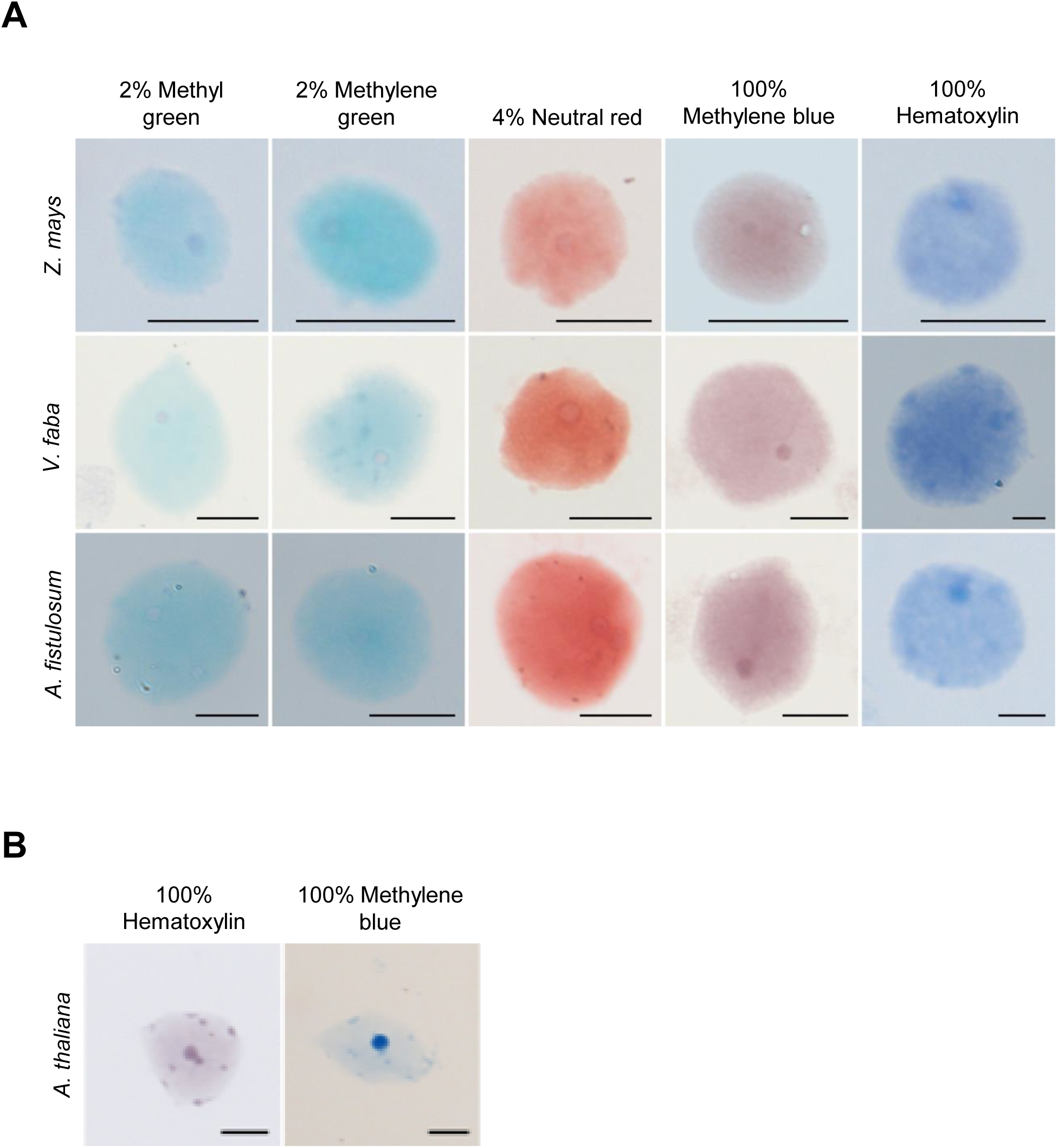
Comparison of different nuclear stains on fixed nuclei. (A) Staining of *Z. mays*, *A. fistulosum* and *V. faba* nuclei with 2% methyl green, 2% methylene green, 4% neutral red, 100% hematoxylin (left) and 100% methylene blue (right) solution for 2 min each. (B) Staining of *A. thaliana* nuclei with 100% hematoxylin (right) and 100% methylene blue (left) solution. Hematoxylin and methylene blue also stained the chromocenters. Scale bars, 5 μm.

**S3 Fig:**
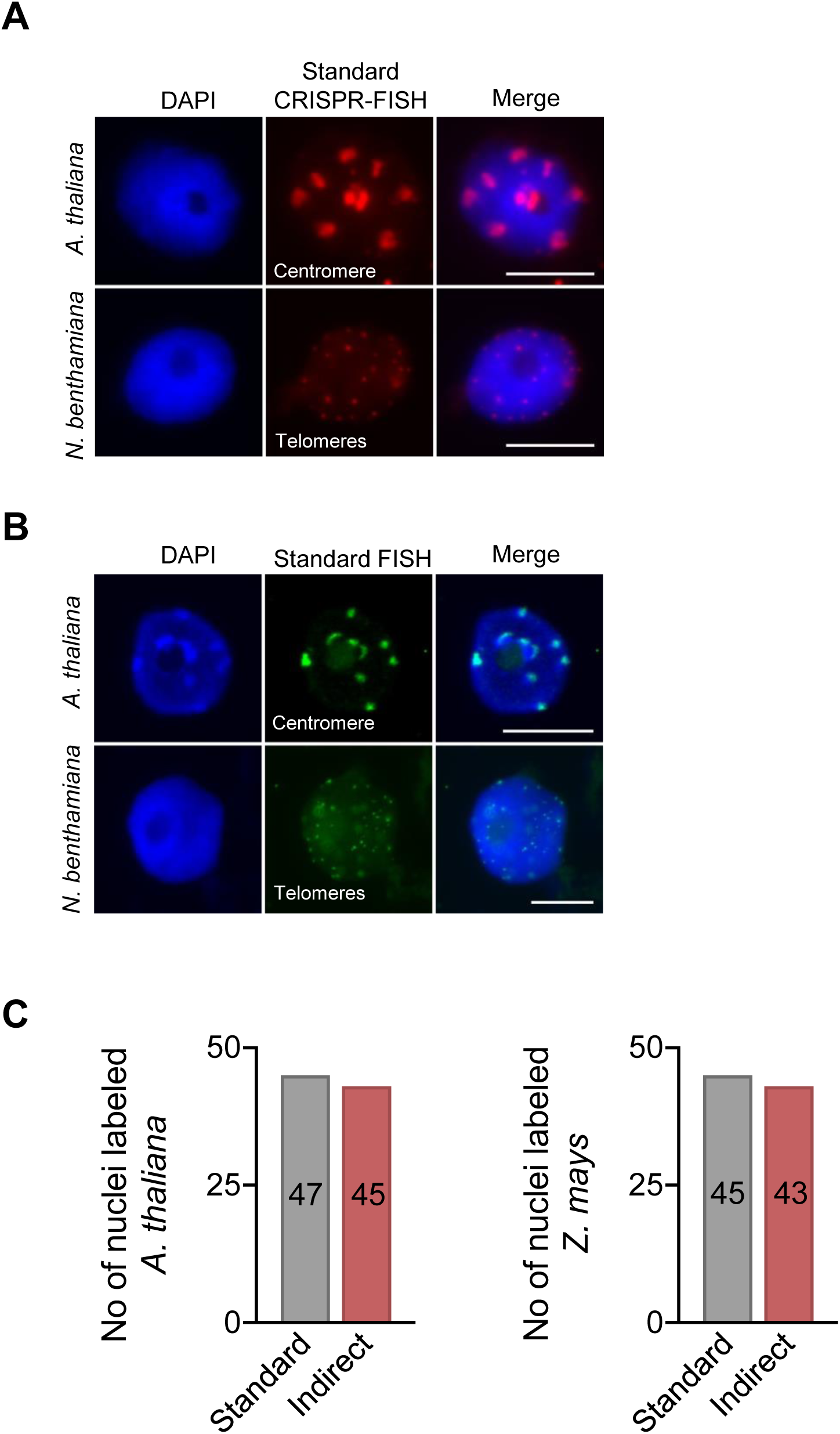
Labeling of DNA repeats using standard CRISPR-FISH and DNA FISH. Visualization of centromere and telomere repeats on formaldehyde fixed *A. thaliana* and *N. benthamiana* with (A) standard CRISPR-FISH (top) using Atto550 tracrRNA and (B) DNA FISH using biotin-labeled oligo probes and detected with streptavidin FITC, respectively. DNA stained with DAPI (blue). Scale bars, 5 μm. (C) Bar graph comparing the efficiency of standard CRISPR-FISH and indirect CRISPR-FISH methods. The number of nuclei with centromere and knob repeats on fixed 2C nuclei of (left) *A. thaliana* and (right) *Z. mays* is shown. n =50 nuclei.

**S4 Fig:**
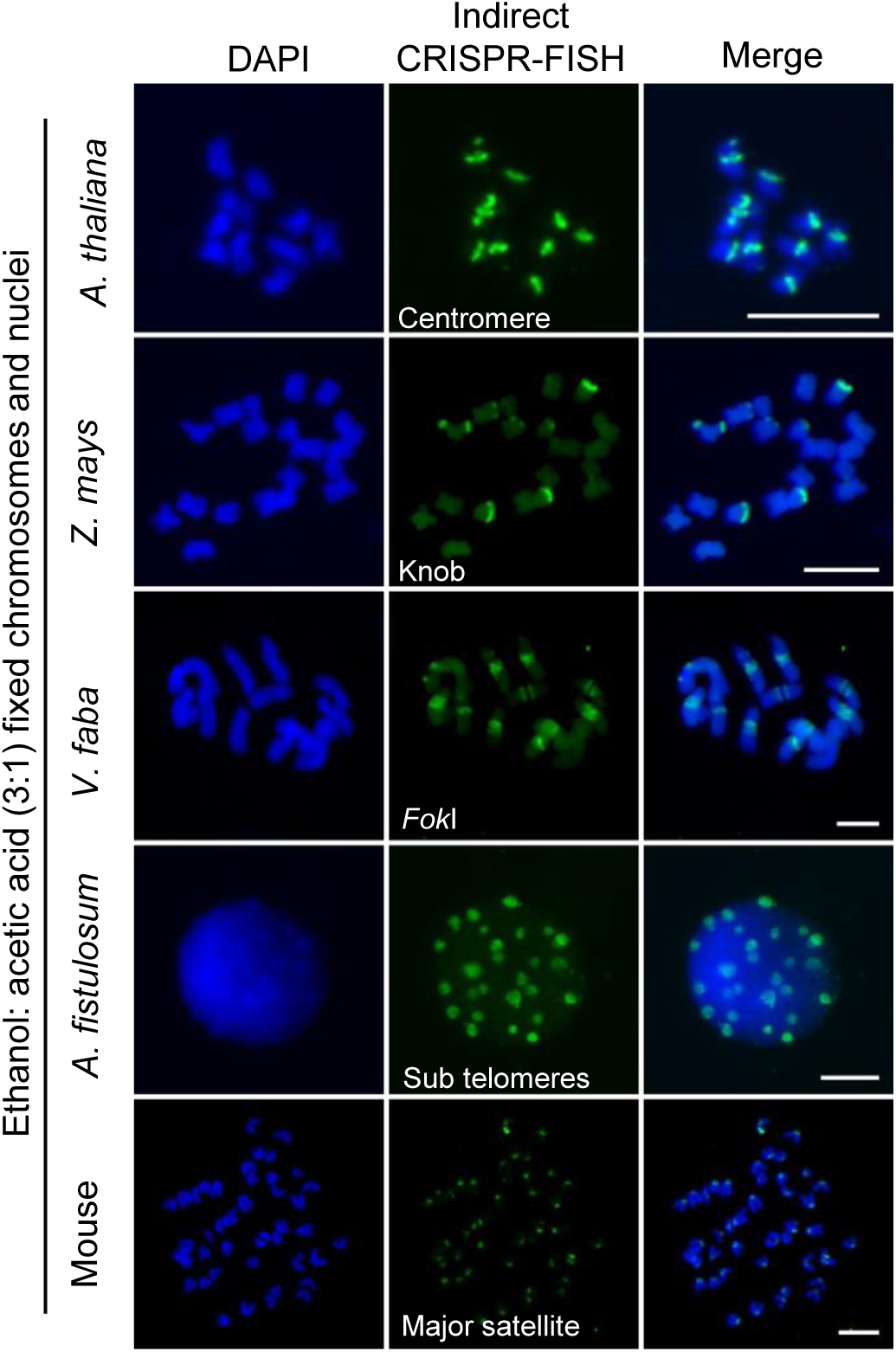
Indirect CRISPR-FISH labeling on ethanol: acetic acid fixed nuclei and chromosomes. Visualization of centromere, knob, *Fok*I, sub telomere and major satellite repeats in ethanol: acetic acid fixed chromosomes of *A. thaliana*, *Z. mays*, *V. faba*, *A. fistulosum* and mouse. Labeling was performed using indirect CRISPR-FISH with biotinylated tracrRNA, followed by detection with streptavidin FITC. DNA stained with DAPI (blue). Scale bars, 5 μm.

**S1 Table:**
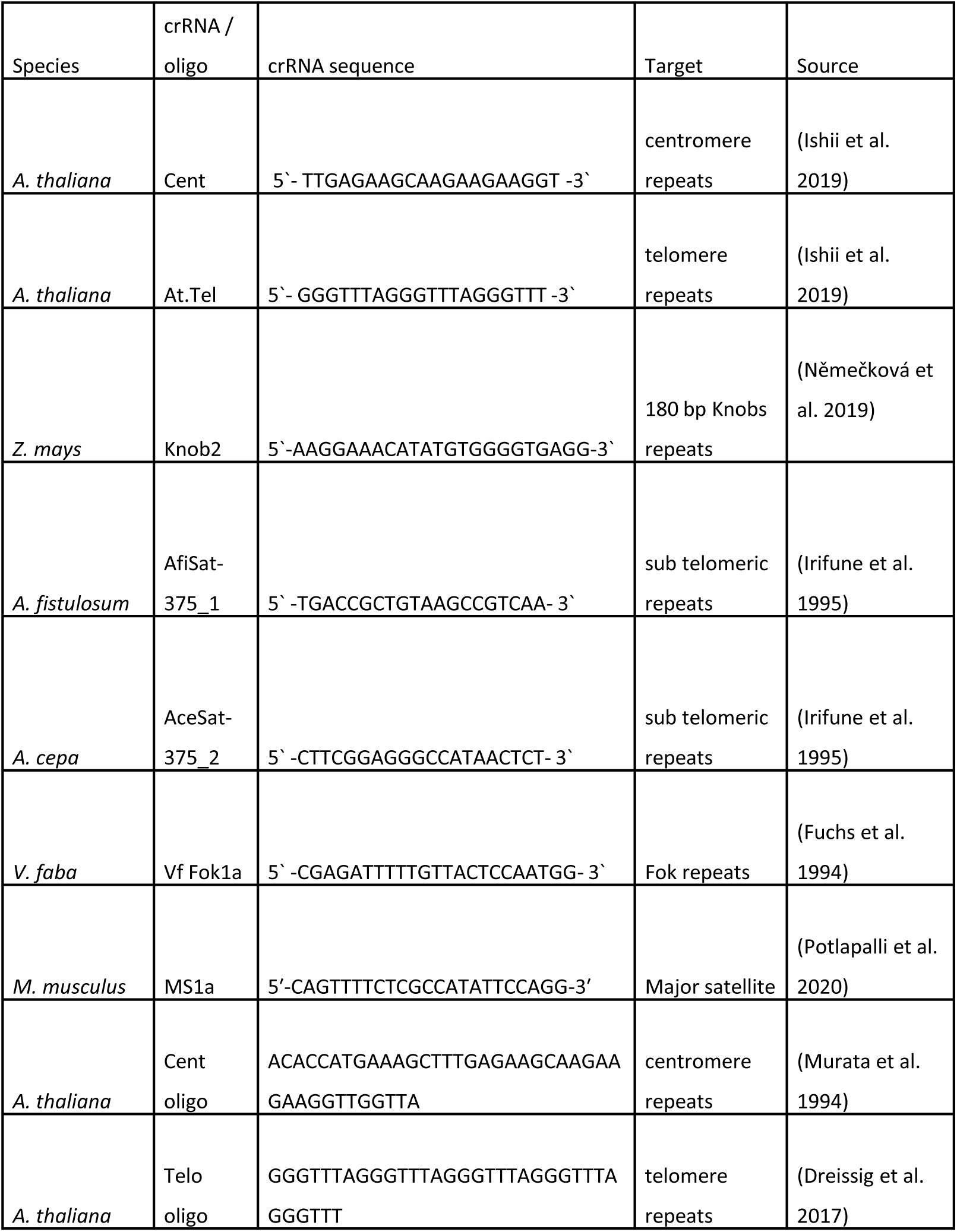
List of crRNA sequences and probe DNA

## References

1. Brigati DJ, Myerson D, Leary JJ, Spalholz B, Travis SZ, Fong CKY, et al. Detection of viral genomes in cultured cells and paraffin-embedded tissue sections using biotin-labeled hybridization probes. Virology. 1983;126(1):32–50.

2. Kozubek S, Lukášová E, Amrichová J, Kozubek M, Lišková A, Šlotová J. Influence of Cell Fixation on Chromatin Topography. Analytical Biochemistry. 2000;282(1):29–38.

3. Shim AR, Frederick J, Pujadas EM, Kuo T, Ye IC, Pritchard JA, et al. Formamide denaturation of double-stranded DNA for fluorescence in situ hybridization (FISH) distorts nanoscale chromatin structure. PLoS One. 2024;19(5):e0301000.

4. Ishii T, Schubert V, Khosravi S, Dreissig S, Metje-Sprink J, Sprink T, et al. RNA-guided endonuclease – in situ labelling (RGEN-ISL): a fast CRISPR/Cas9-based method to label genomic sequences in various species. New Phytologist. 2019;222(3):1652–61.

5. Potlapalli BP, Schubert V, Metje-Sprink J, Liehr T, Houben A. Application of Tris-HCl Allows the Specific Labeling of Regularly Prepared Chromosomes by CRISPR-FISH. Cytogenetic and Genome Research. 2020;160(3):156–65.

6. Deng W, Shi X, Tjian R, Lionnet T, Singer RH. CASFISH: CRISPR/Cas9-mediated in situ labeling of genomic loci in fixed cells. Proceedings of the National Academy of Sciences. 2015;112(38):11870–5.

7. Němečková A, Wäsch C, Schubert V, Ishii T, Hřibová E, Houben A. CRISPR/Cas9-Based RGEN-ISL Allows the Simultaneous and Specific Visualization of Proteins, DNA Repeats, and Sites of DNA Replication. Cytogenetic and Genome Research. 2019;159(1):48–53.

8. Nagaki K, Yamaji N. Decrosslinking enables visualization of RNA-guided endonuclease–in situ labeling signals for DNA sequences in plant tissues. Journal of Experimental Botany. 2020;71(6):1792–800.

9. Potlapalli BP, Fuchs J, Rutten T, Meister A, Houben A. The potential of ALFA-tag and tyramide-based fluorescence signal amplification to expand the CRISPR-based DNA imaging toolkit. Journal of Experimental Botany. 2024.

10. Jayakodi M, Golicz AA, Kreplak J, Fechete LI, Angra D, Bednář P, et al. The giant diploid faba genome unlocks variation in a global protein crop. Nature. 2023;615(7953):652–9.

11. Liao N, Hu Z, Miao J, Hu X, Lyu X, Fang H, et al. Chromosome-level genome assembly of bunching onion illuminates genome evolution and flavor formation in Allium crops. Nature Communications. 2022;13(1):6690.

12. Bogen SA, Vani K, Sompuram SR. Molecular mechanisms of antigen retrieval: antigen retrieval reverses steric interference caused by formalin-induced cross-links. Biotechnic & Histochemistry: Official Publication of the Biological Stain Commission. 2009;84(5):207–15.

13. O’Leary T, Fowler C, Evers D, Mason J. Protein fixation and antigen retrieval: chemical studies. Biotechnic & Histochemistry. 2009;84(5):217–21.

14. Kolodziej KE, Pourfarzad F, de Boer E, Krpic S, Grosveld F, Strouboulis J. Optimal use of tandem biotin and V5 tags in ChIP assays. BMC Molecular Biology. 2009;10(1):6.

15. Bradshaw RA, Cancedda F, Ericsson LH, Neumann PA, Piccoli SP, Schlesinger MJ, et al. Amino acid sequence of Escherichia coli alkaline phosphatase. Proceedings of the National Academy of Sciences. 1981;78(6):3473–7.

16. Rennke HG, Venkatachalam MA. Chemical modification of horseradish peroxidase. Preparation and characterization of tracer enzymes with different isoelectric points. Journal of Histochemistry & Cytochemistry. 1979;27(10):1352–3.

17. Cybulski JS, Clements J, Prakash M. Foldscope: Origami-Based Paper Microscope. PLOS ONE. 2014;9(6):e98781.

18. Collias D, Beisel CL. CRISPR technologies and the search for the PAM-free nuclease. Nature Communications. 2021;12(1):555.

19. Potlapalli BP, Ishii T, Nagaki K, Somasundaram S, Houben A. CRISPR-FISH: A CRISPR/Cas9-Based In Situ Labeling Method. Methods Mol Biol. 2023;2672:315–35.

20. Mandáková T, Lysak MA. Chromosome Preparation for Cytogenetic Analyses in Arabidopsis. Current Protocols in Plant Biology. 2016;1(1):43–51.

21. Khosravi S, Schindele P, Gladilin E, Dunemann F, Rutten T, Puchta H, et al. Application of aptamers improves CRISPR-based live imaging of plant telomeres. Frontiers in plant science. 2020:1254.

22. Murata M, Ogura Y, Motoyoshi F. Centromeric repetitive sequences in *Arabidopsis thaliana*. The Japanese Journal of Genetics. 1994;69(4):361–70.

23. Dreissig S, Schiml S, Schindele P, Weiss O, Rutten T, Schubert V, et al. Live-cell CRISPR imaging in plants reveals dynamic telomere movements. The Plant Journal: For Cell and Molecular Biology. 2017;91(4):565–73.

24. Ma L, Wu S-M, Huang J, Ding Y, Pang D-W, Li L. Fluorescence in situ hybridization (FISH) on maize metaphase chromosomes with quantum dot-labeled DNA conjugates. Chromosoma. 2008;117(2):181–7.

25. Ahmadli U, Sandmann M, Fuchs J, Lermontova I. Immunolabeling of Nuclei/Chromosomes in Arabidopsis thaliana. Methods in molecular biology (Clifton, NJ). 2022;2382:19–28.

